# The regulatory function of the AAA4 ATPase domain of cytoplasmic dynein

**DOI:** 10.1101/2020.09.20.305243

**Authors:** Xinglei Liu, Lu Rao, Arne Gennerich

## Abstract

Cytoplasmic dynein is the primary motor for microtubule minus-end-directed transport and is indispensable to eukaryotic cells. Although each motor domain of dynein contains three active AAA+ ATPases (AAA1, 3, and 4), only the functions of AAA1 and 3 are known. Here, we use single-molecule fluorescence and optical tweezers studies to elucidate the role of AAA4 in dynein’s mechanochemical cycle. We demonstrate that AAA4 controls the priming stroke of the motion-generating linker, which connects the dimerizing tail of the motor to the AAA+ ring. Before ATP binds to AAA4, dynein remains incapable of generating motion. However, when AAA4 is bound to ATP, the gating of AAA1 by AAA3 prevails and dynein motion can occur. Thus, AAA1, 3, and 4 work together to regulate dynein function. Our work elucidates an essential role for AAA4 in dynein’s stepping cycle and underscores the complexity and crosstalk among the motor’s multiple AAA+ domains.

## Introduction

The cytoplasmic dynein motor complex (referred to here as dynein) is an intricate and ubiquitous biological nanomachine that performs most minus-end directed movement along cellular microtubules (MTs). Through its interaction with cofactors such as dynactin and lissencephaly-1 (Lis1), dynein performs a vast array of functions including intracellular transport of organelles and signaling complexes^1-6^, nuclear transport during neuronal migration^7-9^, regulation of mitotic spindle length and positioning, and removal of checkpoint proteins from the kinetochore during cell division^10-12^. Thus, it is not surprising that dynein dysfunction has been linked to a growing number of human diseases termed “dyneinopathies”^13-15^ including malformation of cortical development (MCD)^16-20^, spinal muscular atrophy (SMA)^21,22^, SMA with lower extremity predominance (SMALED)^23-26^, and others^18,19,27,28-31^. However, dynein’s molecular mechanism, the regulatory functions of its subunits, and the molecular effects of disease mutations, remain largely unclear.

Dynein is the largest (∼2.5 MDa) and most complex cytoskeletal motor protein. It is comprised of two identical heavy chains (HCs) and several subunits^5^. The dynein HC contains an N-terminal dimerizing tail domain and a C-terminal motor domain (MD) or “head” with six tandem-linked AAA+ ATPase modules (AAA: ATPase associated with various cellular activities) arranged in a ring (AAA1-6) (**Fig. 1a**). Only AAA1, 3, and 4 hydrolyze ATP^32^. Three elongated structures emerge from the AAA ring: A ∼15-nm coiled-coil “stalk” that protrudes from AAA4^33^ and separates dynein’s MT-binding domain (MTBD) from the AAA+ ring^34^, an antiparallel coiled-coil called the buttress^34^ (or strut^35^) that emerges from AAA5 and contacts the stalk, and a ∼10-nm “linker” that extends from AAA1 and connects the AAA+ ring to the tail. The linker undergoes conformational changes^36,37^ that generate unidirectional motion and force, and it also controls the buttress-mediated sliding of the stalk helices to shift between weak and strong MT-binding states^38^ (**Fig. 1a,b**).

**Figure 1.**
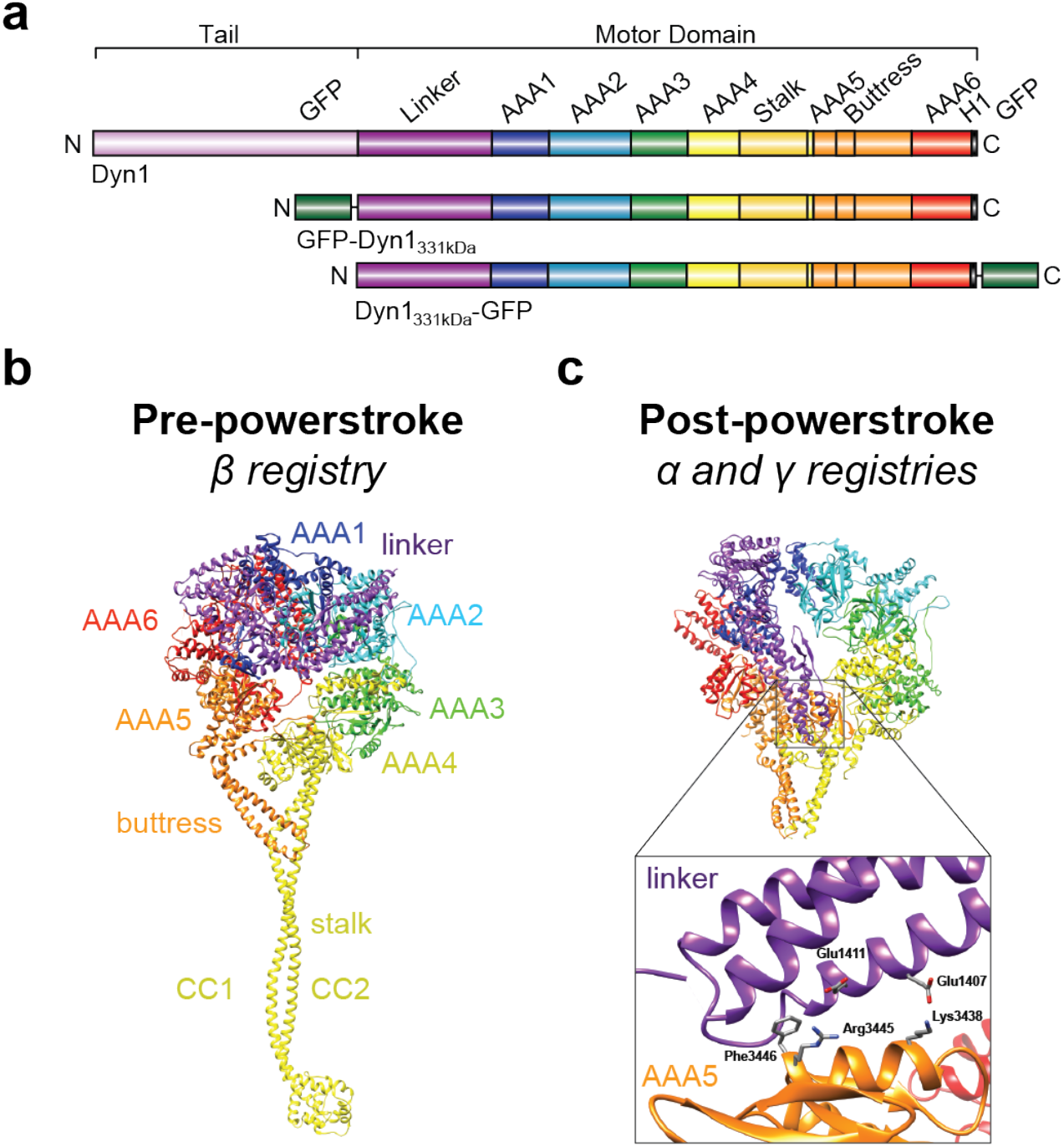
Cytoplasmic dynein domain organization, and the motor domain (MD) pre-and post-powerstroke states in relation to the stalk-helix registrations. (**a**) Organization of the full-length cytoplasmic dynein heavy chain (HC) (a.a. 1–4092) and the tail-truncated monomeric constructs, GFP-Dyn1_331kDa_ and Dyn1_331kDa_-GFP (a.a. 1219–4092). (**b**) Dynein MD structure in the pre-powerstroke state (ADP.Vi, *Homo sapiens* cytoplasmic dynein-2; PDB entry 4RH7^73^). The linker is bent and close to AAA2 (left), and the stalk helices assume the weak microtubule (MT)-binding β registry as a result of the undocked linker^38^. (**c**) Post-powerstroke state (Apo, *S. cerevisiae* dynein; PDB entry 4W8F^33^). The linker is straight and docked on AAA5, and the stalk helices assume the strong MT-binding α registry or the γ registry with intermediate MT-binding strength. Interactions between hydrophobic linker residues (E1407 and E1411) and highly conserved AAA5 residues (F3446, R3445, and K3438) facilitate docking of the linker N-terminus on AAA5^33^ (PDB entry 4W8F^74^) (inset).

Biochemical studies demonstrate that the stalk is a bidirectional communication pathway between the AAA+ ring and MTBD^38-40^. The nucleotide state of the MD affects MT binding and vice versa^41-43^. Furthermore, the stalk-helix registries influence both dynein-MT-binding affinity and MD ATPase activities^38-40^. In solution, the stalk helices predominantly assume a low-affinity registration called the β registration^40,44^. However, after MT binding, the stalk helices transition to the high-affinity α registration^45,46^. In the presence of applied mechanical tension, dynein assumes the strong MT-binding α registry under backward load, while forward load induces the recently discovered γ registry with intermediate MT-binding strength. The stalk assumes the β registry upon AAA1-ATP binding^38^.

Dynein can move processively (the ability to take multiple steps before dissociating), and we are only beginning to understand how dynein generates continuous unidirectional motion under opposing forces. In mammalian dynein, activation by its largest cofactor, dynactin, together with a cargo adaptor (e.g. BICD2) is required to convert dynein from a diffusive^47^/weakly processive^48^ motor to an ultraprocessive motor^49-51^. This activation involves realignment of the MDs into a parallel orientation where both MDs can readily bind the MT^52-54^. Once activated, processive movement can occur under load^55^, with one head binding the MT tightly while the other detaches and advances. Recent studies showed the cargo adaptor-dependent recruitment of two dynein motors arranged side-by-side on a MT (i.e., four MDs in parallel)^56,57^. Recruitment of a second motor likely increases processivity by reducing the dissociation rate of the entire complex from the MT. Although these studies provide important insights into the mechanisms that facilitate dynein processivity, a fundamental question remains: how do the convoluted interactions of dynein’s multiple AAA+ domains control the shifts between strong and weak MT-binding states of dynein’s MTBDs so that the force-bearing leading head stays bound to the MT while the partner head detaches and advances?

It is known that ATP binding to AAA1 and ADP binding to AAA3 both reduce MT-binding strength^58^. Conversely, ADP binding to AAA1 has the opposite effect^58^. In addition, we^58^ and others^59^ reported that AAA3 acts as a “gate” for ATP-induced MT release. ATP binding to AAA1 induces MT release only if AAA3 is in the post-hydrolysis state. When force is applied to the linker, ATP binding to AAA3 “opens” the gate and allows AAA1-mediated, ATP-induced MT release^58^. Adding complexity to the interactions among AAA1, AAA3, dynein MTBDs, and external and intramolecular tension is evidence that AAA4 may have a regulatory role. When ATP hydrolysis by AAA4 is prevented, dynein processivity increases two-fold and MT-binding affinity increases five-fold^60^. However, blockage of AAA4-ATP hydrolysis reduces dynein velocity only slightly^60^, in contrast to prevention of AAA1 or AAA3 ATP hydrolysis, which dramatically effect velocity (prevention of AAA1 hydrolysis makes the motor immobile)^60,61^. It was therefore commonly assumed that AAA4 played a relatively minor role in regulating dynein’s mechanochemical stepping cycle, while AAA1 and AAA3 were the key regulators.

Here, we combine mutagenesis with single-molecule fluorescence and optical tweezers-based force measurements to demonstrate that while blockage of AAA4 ATP hydrolysis reduces the speed of dynein motion only to a small degree, preventing AAA4 ATP binding abolishes dynein motility completely (in contrast to blockage of AAA3 ATP binding), suggesting that AAA4 has an essential regulatory role. We further demonstrate that while AAA3 achieves its effect on dynein-MT binding indirectly via AAA1^58^, AAA4 directly controls the linker conformational change of dynein, and in doing so, governs the buttress-mediated sliding of the stalk helices. We also show that blocking AAA4 ATP binding induces the γ registry independent of the direction of applied tension, and does so irrespective of the nucleotide state of AAA3. However, when AAA4 is bound to ATP, the gating of AAA1 by AAA3 prevails, explaining why the effect of preventing ATP hydrolysis by AAA4 on dynein velocity is so mild. Thus, AAA1, 3, and 4, the three active ATPases of the MD, work together to regulate dynein’s cyclic MT-interactions.

## Results

### AAA4-ATP binding is required for dynein motion

To determine the importance of AAA4 in regulating dynein motion, we used a single-molecule fluorescence motility assay^62^ to define the effects of ATP-binding mutations (K/A mutation in the Walker A motif^36,63^) and ATP-hydrolysis mutations (E/Q mutation in the Walker B motif^63,64^) in AAA3 and AAA4 on dynein motility. We introduced E/Q and K/A mutations in Dyn1_331kDa_, a minimal *S. cerevisiae* MD containing the linker, AAA+ ring, stalk, buttress and MTBD (this construct retains its motor activities^58,65,66^ and is equivalent to the *Dictyostelium discoideum* MD used in key biochemical studies^36,37,39,41,67-69^). We then dimerized the single-headed GFP-Dyn1_331kDa_ mutants using an antibody against the N-terminal GFP^38^ and studied the motors using total internal reflection fluorescence (TIRF) microscopy^62^.

Using this approach, we first confirmed that blocking AAA3-ATP binding or hydrolysis significantly reduced velocities (11.6 ± 0.5 nm/s and 4.8 ± 0.3 nm/s) when compared to WT dynein (110 ± 2 nm/s) (**Fig. 2a**,**b**)^58,59,61,70,71^, while preventing ATP hydrolysis by AAA4 had only minor effects on dynein velocity^60,70^ (velocity reduced to 89 ± 3 nm/s), consistent with previous reports (**Fig. 2a**,**b**). In contrast, blocking ATP binding to AAA4 in the AAA4 K/A single mutant completely abolished dynein motion (as did blocking ATP hydrolysis by both AAA3 and AAA4 in the AAA3 E/Q + AAA4 E/Q double mutant) (**Fig. 2a**). Thus, ATP binding to AAA4 is required for dynein motility.

**Figure 2.**
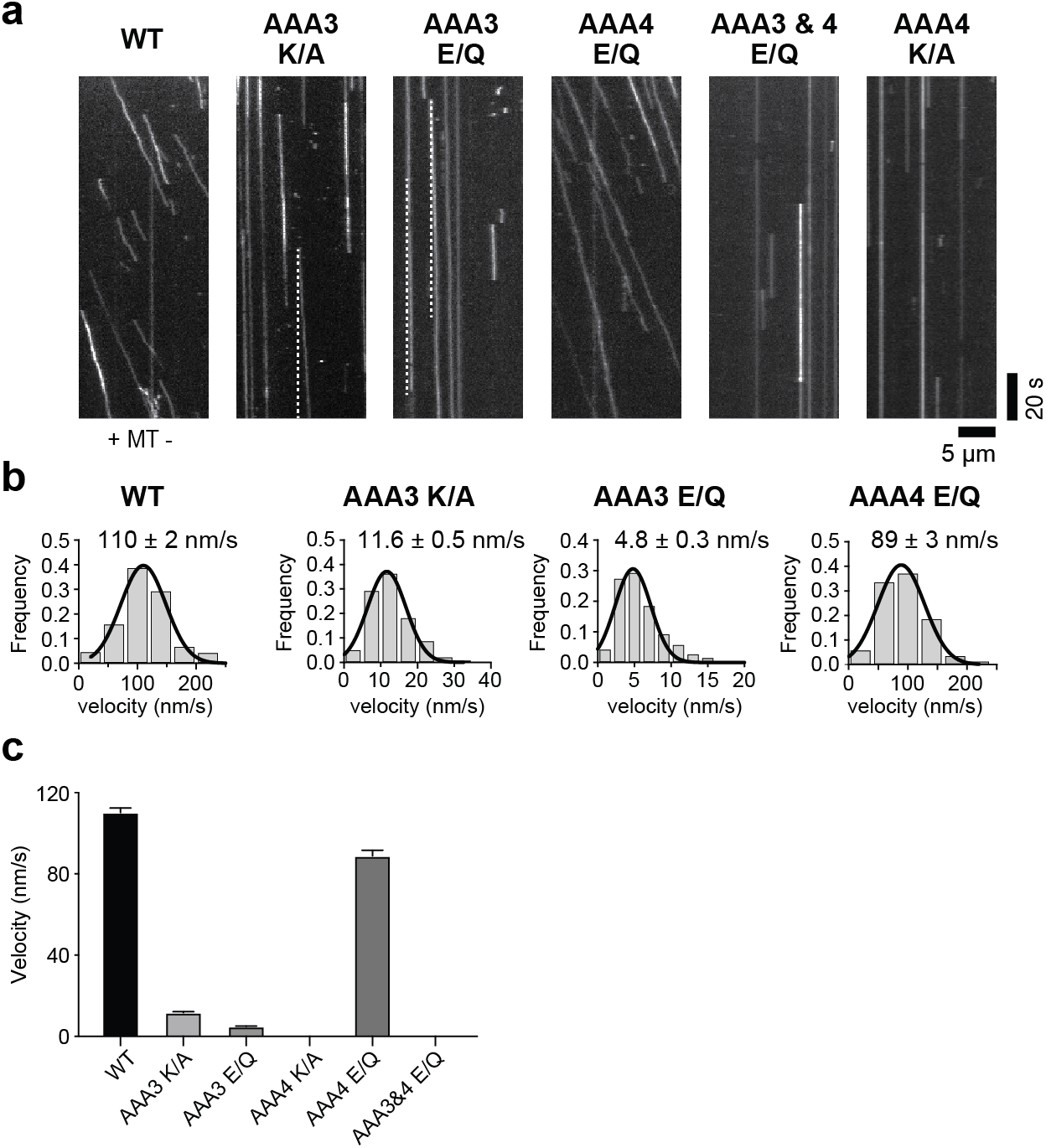
AAA4-ATP binding is essential for dynein motility. (**a**) Antibody-dimerized WT Dyn1_331kDa_ moves processively in the single-molecule TIRF assay. Diagonal lines in the kymograph represent dimerized molecules that are moving over time. Preventing ATP binding (K/A) or ATP hydrolysis (E/Q) by AAA3 dramatically slows down dynein motion (dashed white lines serve as visual guides to identify slow moving mutants). In contrast, preventing ATP hydrolysis by AAA4 only slightly reduces dynein speed. Strikingly, preventing ATP hydrolysis by both AAA3 and AAA4 or preventing AAA4-ATP binding only, completely abolishes dynein motion. (**b**) Fitting the WT velocity histograms with a Gaussian fit (black line), returns a mean velocity of 110 ± 2 nm/s (± SEM; *N* = 284). Analysis of the AAA3 mutants confirms the low velocities suggested by the kymographs: 11.6 ± 0.5 nm/s (± SEM; *N* = 243) for the Dyn1_331kDa_ AAA3 K/A mutant and 4.8 ± 0.3 nm/s (*N* = 295) for the Dyn1_331kDa_ AAA3 E/Q mutant. The Dyn1_331kDa_ AAA4 E/Q mutant moves with a mean velocity of 89 ± 3 nm/s (*N* = 307). (**c**) Comparison of the WT GFP-Dyn1_331kDa_, AAA3 K/A GFP-Dyn1_331kDa_, AAA3 E/Q GFP-Dyn1_331kDa_, and AAA4 E/Q GFP-Dyn1_331kDa_ mean velocities (± SEM).

### ATP binding to AAA4 is required for dynein’s priming stroke

The finding that blocking AAA4-ATP binding completely prevents dynein motion (**Fig. 2a**) surprised us, given the common assumption that AAA4 plays a minor role in regulating dynein stepping relative to AAA1 and AAA3^61,70,72^. We next sought to understand how the AAA4 mutation prevents dynein motion. For dynein to take a forward step, ATP must bind to AAA1, which causes dissociation of the linker from the rear head AAA5 domain^73^ (**Fig. 1b**), transition of the stalk helices from the γ registry into the weak MT-binding β registry^38^, followed by MT release. Linker undocking from AAA5 and subsequent rear head dissociation from the MT are required for the linker to complete its priming stroke^38^. We hypothesized that AAA4 might regulate linker undocking from AAA5, either directly, due to its proximity to the linker docking site, or indirectly, through allosterically regulating AAA1 and/or AAA3 activity.

Our previous works demonstrated that linker undocking from AAA5 is required for the transition from the γ registry into the β registry^38^ (**Fig. 1c**). We therefore tested how the AAA4 K/A and AAA4 E/Q mutations affected stalk registry under load. We performed optical tweezers-based unbinding-force experiments using Dyn1_331kDa_ bearing a C-terminal GFP (**Fig. 1a**) for the attachment to optical trapping beads as we have done before^38,58^ (**Fig. 3a**). Dyn1_331kDa_-GFP contains the linker, AAA+ ring, stalk, buttress and MTBD (**Fig. 1a**), and the linker of this construct can freely assume the pre-and post-powerstroke states in the absence of applied tension (**Fig. 1b**,**c**). Under C-terminus-applied tension, ATP binding to AAA1 causes dissociation of the linker and the expected transition of the stalk helices from the γ to the β registry^38^.

**Figure 3.**
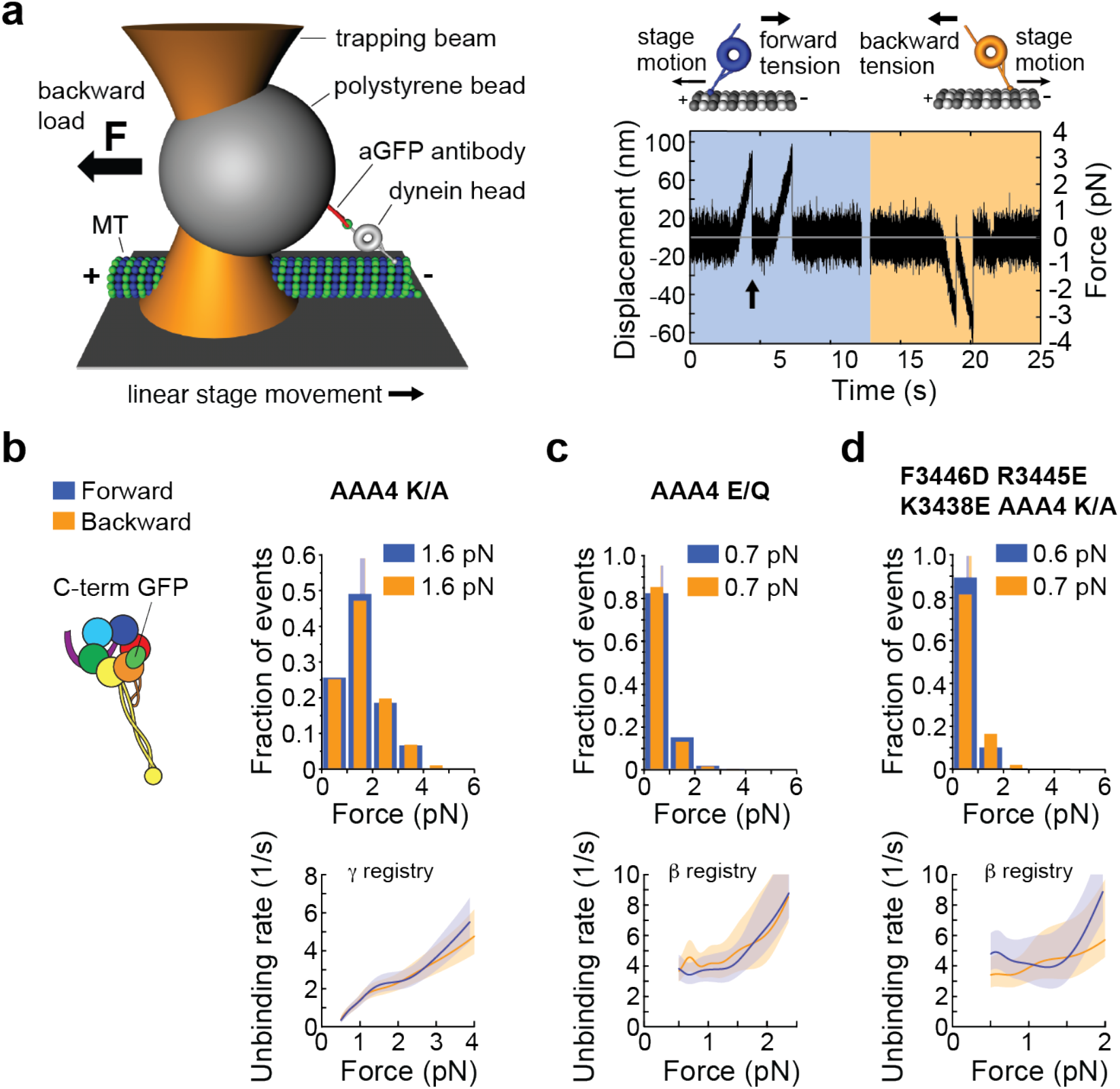
Under C-terminal tension, AAA4-ATP binding controls the linker undocking and the transition from the γ registry with intermediate MT-binding strength to the weak MT-binding β registry. (**a**) (Left) A polystyrene bead bearing a dynein motor is held in an optical trap as the microscope stage sweeps back and forth parallel to a MT (not to scale). (Right) Position (force) vs. time for the AAA4 K/A Dyn1_331kDa_-GFP mutant in the presence of 1 mM ATP. Orange and blue shaded areas show periods of applied backward and forward tension, respectively (loading rate: 5.6 pN/s; k = 0.036 pN/nm, v_stage_ = 156 nm/s). After the motor binds the MT, it pulls the bead out of the trap. Force on the motor increases until the dynein-MT bond ruptures at the “unbinding force” (arrow), here ∼3 pN. (**b**) (Left) Schematic of dynein with GFP fused to the C-terminus. (Right) Normalized histograms of primary forward and backward unbinding forces of the AAA4 K/A Dyn1_331kDa_-GFP mutant in the presence of 1 mM ATP. The mean values are noted. Tall vertical bands represent 95% CIs of the means (forward: [1.5, 1.7] pN, backward: [1.5, 1.7] pN), which were estimated by bootstrapping 4,000 samples. (Bottom) Unbinding rate vs. force derived from the data above. The shaded areas are 95% CIs for the mean rates, which were also estimated by bootstrapping. (**c**) Same as *b*, but for the AAA4 E/Q mutant (95% CIs [0.64, 0.70] and [0.64, 0.71] pN). (**d**) Same as in *b*, but for the F3446D R3445E K3438E AAA4 K/A-Dyn1_331kDa_-GFP mutant (95% CIs [0.63, 0.7] and [0.65, 0.72] pN). The number of events in the forward and backward directions: (*b*) (269, 278), (*c*) (409, 429), and (*d*) (236, 151).

We determined the MT-binding strengths of both AAA4 mutants under C-terminus applied tension and compared their behavior with that of WT Dyn1_331kDa_ with the stalk helices cross-linked (CL) into different registries (we previously generated constructs with cysteines introduced in the outgoing (CC1) and return (CC2) α-helices of the dynein stalk (**Fig. 1b**) for reversible disulfide cross-linking, which locks Dyn1_331kDa_ into fixed stalk registries^38^). In the presence of 1 mM ATP, the AAA4 K/A mutant with the C-terminal GFP exhibited indistinguishable behavior from the Dyn1_331kDa_ mutant with the cross-linked γ registry, Dyn1_331kDa_-γ CL (**Fig. 3b and Table 1**). In contrast, the AAA4 E/Q mutant in the presence of 1 mM ATP behaved similar to Dyn1_331kDa_-β CL, the Dyn1_331kDa_ mutant with the cross-linked β registry (**Fig. 3c and Table 1**). These results indicate that ATP binding to AAA4 is required for linker dissociation from AAA5 and subsequent transition of the stalk helices into the β-registry.

**Table 1.**

| **Experiment 1 (pN),**<br>**mean [CI]** | **Experiment 2 (pN),**<br>**mean [CI]** | **$p_m$** |
| --- | --- | --- |
| AAA4 K/A Dyn1<sub>331kDa</sub>-GFP 1 mM<br>ATP forward<br>1.6 [1.5, 1.7] | Dyn1<sub>331kDa</sub>-γ CL apo forward<sup>38</sup>,<br>1.6 [1.5, 1.7] | < 0.75 |
| AAA4 K/A Dyn1<sub>331kDa</sub>-GFP 1 mM<br>ATP backward<br>1.6 [1.5, 1.7] | Dyn1<sub>331kDa</sub>-γ CL apo backward<sup>38</sup><br>1.6 [1.6, 1.7] | < 0.65 |
| AAA4 E/Q Dyn1<sub>331kDa</sub>-GFP 1 mM<br>ATP forward<br>0.7 [0.6, 0.7] | Dyn1<sub>331kDa</sub>-β CL apo forward<sup>38</sup><br>0.7 [0.6, 0.7] | < 0.2 |
| AAA4 E/Q Dyn1<sub>331kDa</sub>-GFP 1 mM<br>ATP backward<br>0.7 [0.6, 0.7] | Dyn1<sub>331kDa</sub>-β CL apo backward<sup>38</sup><br>0.7 [0.7, 0.8] | < 0.4 |
| F3446D R3445E K3438E AAA4 K/A-Dyn1<sub>331kDa</sub>-GFP 1 mM ATP forward<br>0.6 [0.6, 0.7] | Dyn1<sub>331kDa</sub>-β CL apo forward<sup>38</sup><br>0.7 [0.6, 0.7] | < 0.22 |
| F3446D R3445E K3438E AAA4 K/A-Dyn1<sub>331kDa</sub>-GFP 1 mM ATP backward<br>0.7 [0.7, 0.7] | Dyn1<sub>331kDa</sub>-β CL apo backward<sup>38</sup><br>0.7 [0.7, 0.8] | < 0.45 |
| AAA1 E/Q + AAA4 E/Q-GFP-Dyn1<sub>331kDa</sub> 1 mM ATP forward<br>0.7 [0.6, 0.7] | Dyn1<sub>331kDa</sub>-β CL apo forward<sup>38</sup><br>0.7 [0.6, 0.7] | < 0.83 |
| AAA1 E/Q + AAA4 E/Q-GFP-Dyn1<sub>331kDa</sub> 1 mM ATP backward<br>0.7 [0.6, 0.7] | Dyn1<sub>331kDa</sub>-β CL apo backward<sup>38</sup><br>0.7 [0.7, 0.8] | < 0.38 |
| AAA1 E/Q + AAA3 E/Q + AAA4<br>E/Q-GFP-Dyn1<sub>331kDa</sub> 1 mM ATP<br>forward<br>0.7 [0.6, 0.7] | Dyn1<sub>331kDa</sub>-β CL apo forward<sup>38</sup><br>0.7 [0.6, 0.7] | < 0.83 |
| AAA1 E/Q + AAA3 E/Q + AAA4<br>E/Q-GFP-Dyn1<sub>331kDa</sub> 1 mM ATP<br>backward<br>0.7 [0.6, 0.7] | Dyn1<sub>331kDa</sub>-β CL apo backward<sup>38</sup><br>0.7 [0.7, 0.8] | < 0.3 |
| AAA1 E/Q + AAA4 K/A-GFP-Dyn1<sub>331kDa</sub> 1 mM ATP forward<br>1.6 [1.4, 1.7] | Dyn1<sub>331kDa</sub>-γ CL apo forward<sup>38</sup><br>1.6 [1.5, 1.7] | < 0.96 |
| AAA1 E/Q + AAA4 K/A-GFP-Dyn1<sub>331kDa</sub> 1 mM ATP backward<br>1.7 [1.5, 1.9] | Dyn1<sub>331kDa</sub>-γ CL apo backward<sup>38</sup><br>1.6 [1.6, 1.7] | < 0.66 |
| AAA1 E/Q + AAA3 E/Q + AAA4<br>K/A-GFP-Dyn1<sub>331kDa</sub> 1 mM ATP<br>forward<br>1.5 [1.4, 1.7] | Dyn1<sub>331kDa</sub>-γ CL apo forward<sup>38</sup><br>1.6 [1.5, 1.7] | < 0.62 |
| AAA1 E/Q + AAA3 E/Q + AAA4 K/A-GFP-Dyn1<sub>331kDa</sub> 1 mM ATP backward<br>1.6 [1.5, 1.7] | Dyn1<sub>331kDa</sub>-γ CL apo backward<sup>38</sup><br>1.6 [1.6, 1.7] | < 0.33 |
| AAA1 E/Q + AAA3 K/A + AAA4 K/A-GFP-Dyn1<sub>331kDa</sub> 1 mM ATP forward<br>1.5 [1.4, 1.6] | Dyn1<sub>331kDa</sub>-γ CL apo forward<sup>38</sup><br>1.6 [1.5, 1.7] | < 0.32 |
| AAA1 E/Q + AAA3 K/A + AAA4 K/A-GFP-Dyn1<sub>331kDa</sub> 1 mM ATP backward<br>1.5 [1.5, 1.7] | Dyn1<sub>331kDa</sub>-γ CL apo backward<sup>38</sup><br>1.6 [1.6, 1.7] | < 0.53 |
| AAA3 E/Q + AAA4 E/Q-Dyn1<sub>331kDa</sub>-GFP 1 mM ATP forward<br>1.6 [1.5, 1.7] | Dyn1<sub>331kDa</sub>-γ CL apo forward<sup>38</sup><br>1.6 [1.5, 1.7] | < 0.8 |
| AAA3 E/Q + AAA4 E/Q-Dyn1<sub>331kDa</sub>-GFP 1 mM ATP backward<br>2.6 [2.4, 2.9] | Dyn1<sub>331kDa</sub>-α CL apo backward<sup>38</sup><br>[2.5, 3] | < 0.7 |

### Blocking AAA4-ATP binding prevents sliding of the stalk helices

Interestingly, blocking AAA4-ATP binding induced the γ registry in both MT directions (**Fig. 3a**,**b**), preventing the motor from assuming the strong MT-binding α-registry under backward load. This is a surprising finding as WT dynein in the absence of ATP (apo) and the AAA1 K/A single mutant in the presence of ATP both assume the α registry under backward load^38,58^. The finding that the AAA4 K/A mutant assumes the γ registry under backward load therefore demonstrates unexpectedly that the sliding of the stalk helices is prevented when AAA4-ATP binding is blocked.

As the linker must be docked to AAA5 for the α and γ registries^38^ and since the linker is physically close to AAA4 in the post-powerstroke state^33^ (**Fig. 1c**), we speculated that AAA4 acted directly on the linker to prevent stalk-helix sliding and to lock in the γ registry in both directions. To provide evidence for this possibility, in the AAA4 K/A mutant background with the C-terminal GFP, we mutated three conserved residues in the AAA5 linker-docking site (F3446D, R3445E, and K3438E) that have been previously shown to be critical for the docking of the linker to AAA5^33^ (**Fig. 1c**). We anticipated that this mutant would assume the β registry despite having AAA4-ATP binding blocked. Indeed, the F3446D R3445E K3438E AAA4 K/A Dyn1_331kDa_ quadruple mutant exhibited statistically indistinguishable behavior from the Dyn1_331kDa_-β CL mutant (**Fig. 3d and Table 1**). This result reveals that linker-AAA5 interactions play an important role in AAA4’s ability to prevent the sliding of the stalk helices and to lock in the γ registration.

### AAA4-based regulation of linker undocking is independent of AAA3

Previous work indicates that AAA3 gates AAA1 activity so that AAA1-ATP binding induces the undocking of the linker and the weak MT-binding β registration only when AAA3 is in the post-hydrolysis state^58,59^. However, when tension is applied to the linker, ATP binding to AAA3 is sufficient to “open” the gate^58^. The conclusion that AAA3 affects the linker indirectly via AAA1 is supported by the MT-binding behavior of the AAA3 E/Q mutant. Blocking AAA3-ATP hydrolysis yields the γ registry under forward load and the α registry under backward load^38,58^. This is the same behavior as the AAA1-ATP binding (AAA1 K/A) mutant and the WT motor in the apo state ^38,48^ discussed above. Thus, while AAA3 regulates linker docking and stalk registry indirectly via AAA1, our results suggest that the nucleotide state of AAA4 directly effects the linker.

To provide further evidence that AAA4 controls the docking of the linker directly and to elucidate the hierarchy between the AAA4-induced block on the linker transition from a docked to an undocked state and AAA3 function, we employed novel double and triple AAA+ ATPase mutants. Under linker-applied tension, the binding of ATP to AAA3 opens the gate and AAA1-ATP binding induces the β registry^38,58^. Thus, in the presence of 1 mM ATP, the AAA1 E/Q single mutant and the AAA1 E/Q + AAA3 E/Q double mutant assume the β registry in both directions, while the AAA1 K/A single mutant and the AAA1 E/Q + AAA3 K/A double mutant assume the γ registry under forward load and the α registry under backward load^38,58^. We anticipated that the AAA1 E/Q + AAA4 E/Q double mutant and the AAA1 E/Q + AAA3 E/Q + AAA4 E/Q triple mutant would also assume the β registry in the presence of 1 mM ATP. Additionally, we predicted that blocking AAA4-ATP binding in the double mutant (AAA1 E/Q + AAA4 K/A) and the triple mutant (AAA1 E/Q + AAA3 E/Q + AAA4 K/A) would prevent reduced MT binding and induce the γ registry in both directions. In support of these predictions, the AAA1 E/Q + AAA4 E/Q double mutant and the AAA1 E/Q + AAA3 E/Q + AAA4 E/Q triple mutant assumed the β registry in the presence of 1 mM ATP (**Fig. 4a**,**b and Table 1**), while the AAA1 E/Q + AAA4 K/A double mutant and the AAA1 E/Q + AAA3 E/Q + AAA4 K/A triple mutant assumed the γ registry in both directions (**Fig. 4c**,**d and Table 1**).

**Figure 4.**
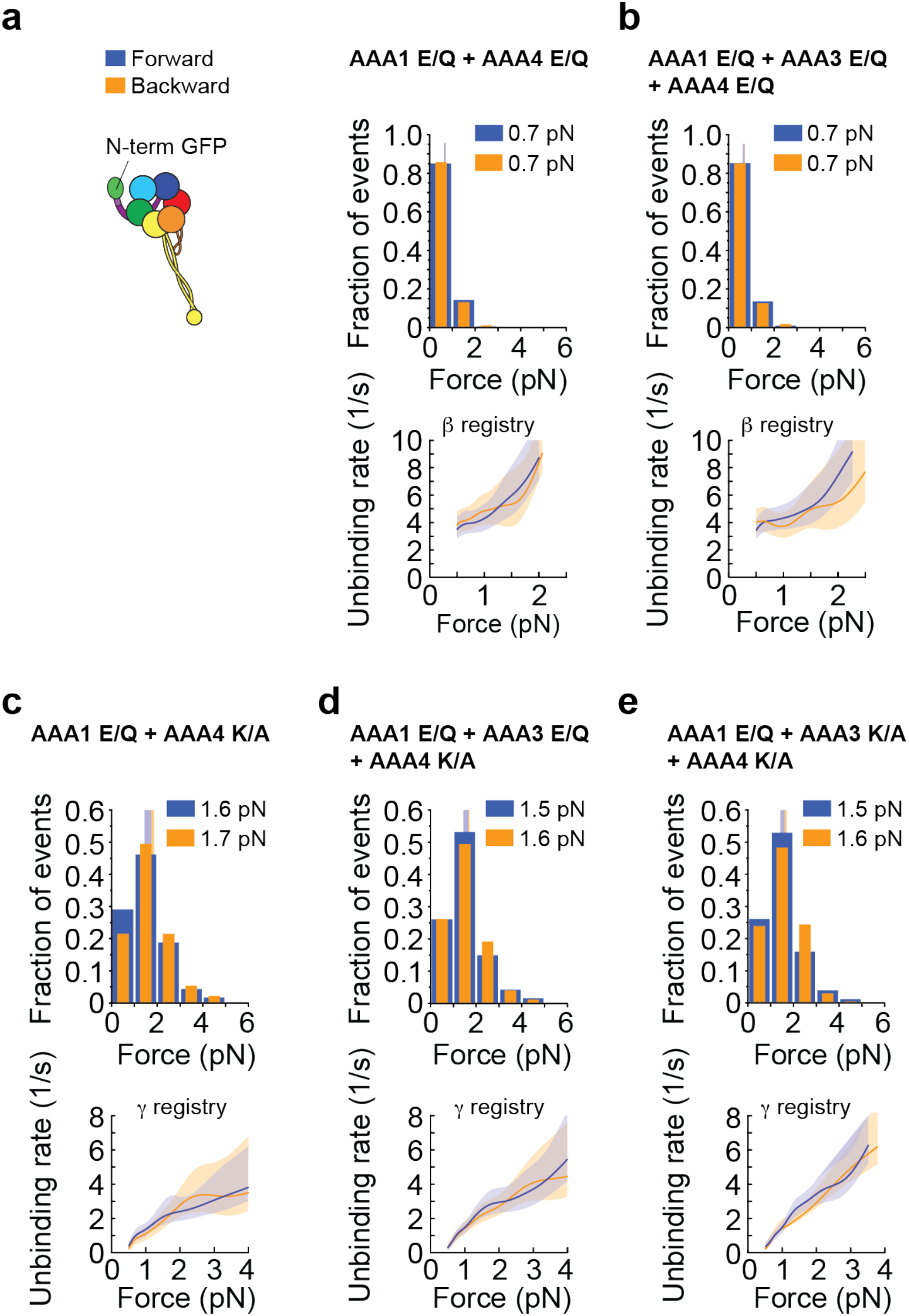
Blocking AAA4-ATP binding induces the γ registry irrespective of the AAA3 nucleotide state. Tension is applied to the N-terminus in the presence of 1 mM ATP. (**a**) (Left) Schematic of dynein with GFP fused to the N-terminus. (Right) Histogram of forward (blue) and backward (orange) unbinding forces for the AAA1 E/Q + AAA4 E/Q-GFP-Dyn1_331kDa_ double mutant measured in the presence of 1 mM ATP. The respective mean values are noted. The vertical bands represent 95% CIs for the means (forward: [0.6, 0.7] pN, backward [0.6, 0.7] pN). (Bottom) Unbinding rate vs. force derived from the data above. The shaded areas are the 95% CIs for the mean rates, which were estimated by bootstrapping. (**b**) Same as in *a*, but for the AAA1 E/Q + AAA3 E/Q + AAA4 E/Q-GFP-Dyn1_331kDa_ triple mutant (95% CIs [0.6, 0.7] and [0.6, 0.7] pN). (**c**) Same as in *a*, but for the AAA1 E/Q + AAA4 K/A-GFP-Dyn1_331kDa_ double mutant (95% CIs [1.4, 1.7] and [1.5, 1.9] pN). (**d**) Same as in *a*, but for the AAA1 E/Q + AAA3 E/Q + AAA4 K/A-GFP-Dyn1_331kDa_ triple mutant (95% CIs [1.4, 1.7] and [1.5, 1.7] pN). (**e**) Same as in *a*, but for the AAA1 E/Q + AAA3 K/A + AAA4 K/A-GFP-Dyn1_331kDa_ triple mutant (95% CIs [1.4, 1.6] and [1.5, 1.7] pN). The number of events in the forward and backward directions: (*a*) (321,358), (*b*) (419, 382), (*c*) (117, 93), (*d*) (188,172), and (*e*) (257, 205).

To test whether AAA4 was able to induce the γ registry both when AAA3 was bound to ATP (regulatory gate on AAA1 is open) and when AAA3 was nucleotide-free (gate on AAA1 is closed), we explored the MT-binding behavior of the AAA1 E/Q + AAA3 K/A + AAA4 K/A triple mutant. As expected, this triple mutant also assumed the γ registry in the presence of 1 mM ATP in both directions (**Fig. 4e and Table 1**), which further supported the premise that AAA4 has a dominant role.

In addition, the fact that the AAA1 E/Q + AAA4 K/A double mutant assumed the γ registry irrespective of the direction of applied tension (**Fig. 4d**) demonstrates that the AAA4 K/A mutation also blocked linker undocking even when AAA3 assumed a post-ATP-hydrolysis state (AAA3 of this mutant can bind and hydrolyze ATP and reach the ADP⋅Pi transition state for some fraction of the dynein cycle^59^, which is sufficient to open the gate on AAA1 so that the AAA1 E/Q single mutant assumes the β registry at 1 mM ATP^38,58^). Thus, AAA4’s regulation of linker undocking is independent of the nucleotide state of AAA3.

### AAA3 controls ATP-induced, AAA1-mediated MT release when AAA4 is bound to ATP

Our work revealed that blocking AAA4-ATP binding had a dominant effect over AAA3 function with respect to linker conformation and stalk registration. We next investigated whether AAA4 in a different nucleotide state had a dominant role throughout dynein’s mechanochemical cycle or whether AAA3 could assume its role in controlling AAA1 function^58,59^. To test AAA3 functionality when AAA4 was bound to ATP, we determined the unbinding behavior of the AAA3 E/Q + AAA4 E/Q double mutant with the C-terminal GFP. Under C-terminus-applied tension, blocking AAA3-ATP hydrolysis closes the gate on AAA1 (AAA1-ATP binding cannot induce MT release) and results in the γ registry under forward load and the α registry under backward load^38,58^. In support of a dominant role for AAA4, we anticipated that AAA4-ATP binding would cause linker detachment and the transition into the β registration irrespective of the AAA3-induced block on AAA1. However, we found that the AAA3 E/Q + AAA4 E/Q double mutant in the presence of 1 mM ATP assumed the γ registry under forward load and the α registry under backward load (**Fig. 5 and Table 1**). Thus, when AAA4 was ATP bound, AAA3 prevailed in gating the activity of AAA1. In conclusion, both regulatory domains exert dominant effects on dynein-MT binding that depend on their nucleotide states. This dual dominance underscores the intricate complexity and convoluted interactions among dynein’s multiple AAA+ domains.

**Figure 5.**
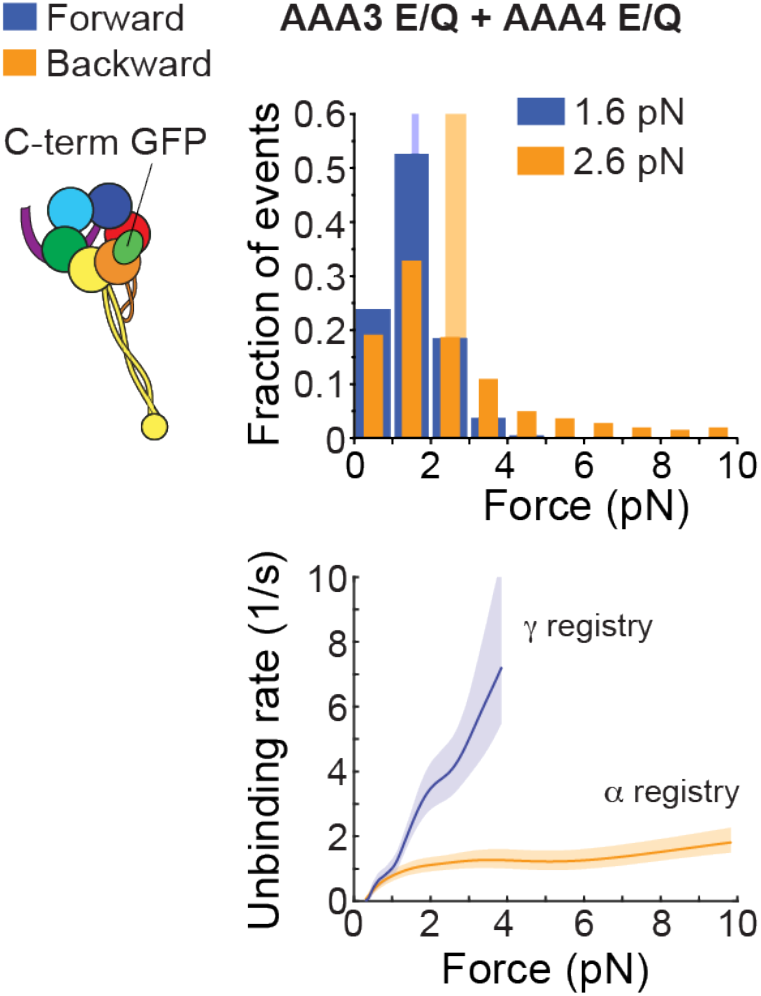
ATP binding to AAA4 is not sufficient to overcome the AAA3-induced block on AAA1. Tension is applied to the C-terminus in the presence of 1 mM ATP. (Left) Schematic of dynein with GFP fused to the C-terminus. (Middle) Histogram of forward (blue) and backward (orange) unbinding forces for the AAA3 E/Q + AAA4 E/Q-Dyn1_331kDa_-GFP mutant measured in the presence of 1 mM ATP. The respective mean values are noted. The vertical bands represent 95% CIs for the means (forward: [1.5, 1.7] pN, backward [2.4, 2.9] pN). The number of events in the forward and backward directions: (279, 234). (Right) Unbinding rate vs. force derived from the data. The shaded areas are 95% CIs for the mean rates, which were estimated by bootstrapping.

## Discussion

Previous studies suggested that AAA4 has only a minor role in dynein function. Here, we show that AAA4 gates the priming stroke of the linker and in this role, it also controls stalk-helix registrations to mediate changes in MT-binding strength. Because gating by AAA4 occurs irrespective of the nucleotide state of AAA3, our results suggest that AAA4 has a major regulatory role in dynein’s stepping cycle, and that its regulation of the linker transition from the post-powerstroke to pre-power stroke state is independent of AAA3 activity. Mutations in AAA4 that cause malformation of cortical development reinforce the importance of AAA4 in dynein mechanochemistry^20^.

Preventing ATP hydrolysis by AAA4 increases dynein processivity two-fold and MT-binding affinity five-fold^60^. However, blocking ATP hydrolysis by AAA4 only reduces dynein velocity by ∼20%^60^ (**Fig. 2**). In contrast, preventing ATP hydrolysis by AAA3 reduces dynein velocity by ∼95%^60,71^ (**Fig. 2**). These initial studies indicated that AAA4 only had a minor regulatory function. However, by combining mutagenesis, single-molecule fluorescence and optical tweezers studies, we reveal that AAA4 has a significant role. Blocking AAA4-ATP binding completely abolished dynein motion and AAA4-ATP binding facilitated the dynein priming stroke. We recently reported that the linker controls the buttress-mediated sliding of the stalk helices to alter the MT-binding strength of the two MTBDs within dynein^38^. These past results combined with the data presented here suggest that AAA4 has an important function in controlling dynein’s cyclic MT interactions. This conclusion appears to contradict a previous study on *Dictyostelium* dynein where it was reported that the AAA4 K/A mutation reduced MT-gliding activity by only 60%^61^. While it is possible that *Dictyostelium and S. cerevisiae* dynein behave differently in the mutant background, future studies will be required to resolve this discrepancy.

Our single-molecule TIRF studies revealed that the AAA3 E/Q + AAA4 E/Q double mutant does not move (**Fig. 2**), which can be explained by the MT-binding behavior of the mutant. Our unbinding-force studies showed that under C-terminus applied tension, AAA3 E/Q + AAA4 E/Q-Dyn1_331kDa_-GFP assumes the γ registry under forward load and the α registry under backward load in the presence of 1 mM ATP (**Fig. 5**). Because both registrations can only be assumed when the linker is docked, the data suggest that the linker is always docked and that the priming stroke never occurs, and thus the motor cannot move. The Dyn1_331kDa_-α CL mutant still moves slowly^38^, which excludes the possibility that the ATP-insensitive MT-binding strength of the AAA3 E/Q + AAA4 E/Q double mutant accounts for its immobility. Intriguingly, the linker of the AAA3 E/Q mutant has also been shown locked in the post-powerstroke state in the absence of a load^74^, but the dimerized AAA3 E/Q mutant moves slowly (**Fig. 2a**,**b**). This implies that the linker of the AAA3 E/Q mutant can detach under tension and undergo a priming stroke, albeit less efficiently whereas the AAA3 E/Q + AAA4 E/Q double mutant cannot. Therefore, the interaction strength between the linker and AAA5 could be stronger when ATP hydrolysis is prevented in both regulatory ATPase domains. The observation that the dimerized AAA4 K/A mutant is immotile further supports the premise that blocking AAA4-ATP binding could result in a stabilized, docked linker conformation.

The finding that blocked AAA4-ATP binding induced the γ registry independent of both the direction of applied tension and the nucleotide state of AAA3 is intriguing because it suggests that AAA4 can prevent tension-and nucleotide-induced changes in the stalk helices. Our data with the F3446D R3445E K3438E AAA4 K/A quadruple mutant showed that the effects of AAA4 on the stalk registry required functional interactions between the linker and AAA5 (**Fig. 3d**). However, our previous work has also demonstrated that the α registry requires the linker to dock to AAA5 in the apo state and that despite the docked state, directional tension can shift the registries interchangeably between α and γ^58^. Blocking AAA4-ATP binding locks the γ registry in both directions and prevents a change to the α registry under backward load, implying that the linker docks to AAA5 in a way that differs from the docked state required for the α registry. Indeed, the crystal structures in the absence of a nucleotide^33^ (**Fig. 1b**) and in the presence of ADP^75^ reveal two distinct post-powerstroke positions. In the apo state, the linker is positioned above the large domain of AAA5 (AAA5L)^33^. In the ADP state, the linker is positioned between AAA5L and the large domain of AAA4 (AAA4L)^75^. Direct linker interaction with AAA4, especially when AAA1 is bound to ATP and AAA4 is nucleotide-free, (**Fig. 4d**), remains to be shown. Solving the crystal structure of the AAA1 E/Q + AAA4 K/A double mutant (**Fig. 4d**) or the AAA1 E/Q + AAA3 E/Q + AAA4 K/A triple mutant (**Fig. 4b**) in the presence of ATP would help to evaluate this possibility.

Our results with the AAA3 E/Q + AAA4 E/Q double mutant exemplified the unique bond that dynein forms with a MT under backward load when the stalk helices assume the α registry. In agreement with our previous work^38,58^, the α registry caused dynein to exhibit “slip-ideal” bonding with the MTs for forces up to ∼8 pN. Faster unbinding was observed for backward forces up to ∼2 pN and a constant, force-independent unbinding rate occurred for greater backward forces (**Fig. 5**). Dynein exhibited slip-bonding up to 4 pN when it assumed the γ registry^38^ (**Figs. 3 & 4**). Noting that the movement of full-length *S. cerevisiae* dynein (Dyn1_471kDa_) ceases at 4.5 pN^76^ (“stall force”), these bond behaviors capture the most relevant force range of dynein.

A recent study reported that a single *S. cerevisiae* dynein head exhibited slip-bonding in both MT directions^77^. Although the authors confirmed that the unbinding rate in the backward direction increased more slowly than in the forward direction^38,58,66,78^, the unbinding rates increased exponentially. However, a data comparison reveals that these results only differ from ours for forces above ∼4 pN (**Figs. 3 & 4**). As we have already discussed^38^, the reason for this discrepancy is primarily due to the different unbinding-force assays that were used: while Ezber et al.^77^ used the “oscillatory assay” where the trap rapidly switched between two positions^66^, we used the “constant-pulling assay” where a MT attached to a coverslip moved at a constant velocity past the stationary trapped bead^38,58,79^. Extremely high loading rates up to ∼25,000 pN/s can develop in the oscillation assay^38^; therefore, it is possible that the high loading rates affect the dynein-MT bond and contribute to increased unbinding rates with larger forces. The loading rates in our constant-pulling assay remained below 5 pN/s for forces up to 12 pN^38^. The maximal loading rates increase significantly with increasing force in the oscillatory assay from ∼2,000 pN/s at 1 pN to ∼25,000 pN/s at 6.5 pN^38^. Importantly, the time period when the extreme loading rates are acting on the MTBD-MT bond is close to the timescale of topological protein reorganization^80^.

In summary, we revealed that blocking AAA4-ATP binding locks the dynein linker element in the post-powerstroke state and mediates the transition of the stalk helices into the γ registry. Our data suggests that ATP binding to AAA4 is necessary for the trailing head to undock the linker from AAA4/AAA5, which allows for the transition into the weak MT-binding β registration^38^ when AAA1 is bound to ATP and AAA3 is in the post-hydrolysis state^58^. This suggest that dynein uses a “two-doored”, double “key-lock” mechanism^81^ where the AAA3^58^ and AAA4 nucleotide states control the effects of both AAA1-ATP binding on dynein-MT binding and AAA1-ATP hydrolysis on linker conformational changes. Multi-layered regulation by dynein’s AAA+ domains underscores the complex interactions among AAA1, 3, and 4, dynein MTBDs, and external and intramolecular tension that collectively control dynein mechanochemistry.

## Methods

### Generation of yeast strains

Mutant yeast stains were engineered as described previously^38^. Briefly, the PrimerQuest tool from IDT (Integrated DNA Technologies) was used to design PCR primers, and DNA fragments were generated using KOD Hot Start DNA polymerase (EMD Millipore). Yeast transformation was performed using the Frozen-EZ Yeast Transformation II Kit (Zymo Research), followed by a two-step selection method using either synthetic media with uracil-dropout amino acid mix (SC/URA) or 5-fluorouracil (5-FOA) as the selective agents. All newly engineered and mutated yeast strains were confirmed by PCR and sequencing.

### Yeast growth and protein expression

Yeast cultures and dynein purification were performed as described previously^38^ with the modifications described below. The parent strain used in this work was VY137, which expresses a tail-truncated minimal *S. cerevisiae* MD containing the linker, AAA+ ring, stalk, buttress, and MTBD. This construct retains its motor activities^58,65,66^ and is equivalent to the *Dictyostelium discoideum* MD used in key biochemical studies^36,37,39,41,67-69^. The genotype was *pGal:ZZ:Tev:GFP:HA:D6 MATa; his3-11,15; ura3-1; leu2-3,112; ade2-1; trp1-1; pep4Δ:HIS5; prb1Δ*. Mutants were expressed behind the inducible galactose promotor (*pGAL*). Yeast cells were grown in 5 mL of 2× YPD medium (20 g/L yeast extract, 40 g/L peptone, 4% [w/v] dextrose) overnight, then transferred to 50 mL of YPR solution (2% [w/v] raffinose) and inoculated for 8 hours. For single-headed dynein, expression was induced by growing the cells in 2× YPG medium (4% [w/v] galactose) to a final OD_600_ between 1.5 and 2.5 (∼16 hours). After harvesting by centrifugation at 500 × g for 5 minutes, the pellets were resuspended in 0.2 volumes of ddH_2_O and flash-frozen in liquid nitrogen as small droplets. The cell pellets were stored at -80°C until further use.

### Yeast dynein purification

To purify the dynein, the frozen cell pellet droplets were pulverized using a kitchen coffee grinder that had been pre-chilled with liquid nitrogen. Next, 0.25 volumes of 4× lysis buffer (1× lysis buffer: 30 mM HEPES, 50 mM KAc, 2 mM Mg(Ac)_2_, 1 mM EGTA, 10% glycerol, 1 mM DTT, 0.1 mM Mg-ATP, 0.5 mM Pefabloc, 10 ng/mL Leupeptin, 10 ng/mL Pepstatin A, and 0.2% v/v Triton X-100, pH 7.2) were added to reach a final 1× lysis buffer concentration. The cell lysate was cleared via ultracentrifugation at 80,000 × g for 10 minutes at 4°C. Then, 0.25 mL of IgG sepharose 6 fast flow beads (GE Healthcare) were added to the supernatant and incubated for 1 hour at 4°C while rotating. The dynein-bound IgG beads were then washed with ∼10 bead volumes of 1× lysis buffer, followed by a wash with 10 mL of 1× TEV protease cleavage buffer (30 mM HEPES, 150 mM KAc, 2 mM Mg(Ac)_2_, 1 mM EGTA, 10% glycerol, 0.1 mM Mg-ATP, 0.5 mM Pefabloc, and 0.1% v/v Triton X-100, pH 7.2). The beads were then resuspended in an equal volume of cleavage buffer and 2% v/v of AcTev protease (ThermoFisher Scientific) was added. The mixture was nutated at 4°C for 2 hours to cleave dynein from the IgG beads. The mixture was centrifuged at 1000 × g for 1 minute at 4°C, and the dynein-containing supernatant was then collected, flash-frozen with liquid nitrogen, and stored at -80°C.

### MT binding and release purification of dynein

MT binding and release purification was performed to further purify the dynein and to remove any inactive motors and aggregates. To 50 μL of TEV-released dynein, 0.1 µL of 10 mM paclitaxel and 5 μL of 5 mg/mL paclitaxel-stabilized MTs were added. This solution was then layered onto a 100 μL sucrose cushion (30 mM HEPES, 200 mM KCl, 2 mM MgCl_2_, pH 7.4, 10% v/v glycerol, 25% w/v sucrose, 1 mM DTT, and 20 μM paclitaxel) and centrifuged at 25°C for 10 minutes at 50,000 × g. After removing the supernatant, the pellet was carefully rinsed with 60 μL of wash buffer (30 mM HEPES, 150 mM KCl, 2 mM MgCl_2_, 10% glycerol, 1 mM EGTA, 1 mM DTT, and 20 µM paclitaxel, pH 7.2) and resuspended in 52 μL of wash buffer with 6 mM Mg-ATP. After incubating for 2 minutes at room temperature, the solution was centrifuged for 5 minutes at 50,000 × g. The MT suspension was then collected, aliquoted in 2 μL volumes, and flash-frozen in liquid nitrogen before storing at -80°C.

### Flow chamber preparation and MT immobilization

Flow chambers were prepared as described previously^82^. Briefly, 18 × 18 × 0.17-mm coverslips (Zeiss) were placed in a porcelain coverslip rack using forceps and submerged in HNO_3_ (25% v/v) for 15 minutes followed by rinsing with ddH_2_O. Then, the rack with the coverslips was placed in NaOH for 2 to 5 minutes, followed by rinsing extensively with ddH_2_O. After drying on a heating block for 30 minutes, the coverslips were store in a vacuum desiccator. The flow chambers were assembled as described^8^. To immobilize MTs on the coverslip surface, 10 μL of 5 mg/mL biotinylated α-casein was flowed into the slide chamber and incubated for 10 minutes. The chamber was then washed three times with 20 μL of blocking buffer (“BB”, 80 mM PIPES, 2 mM MgCl_2_, 1 mM EGTA, 1% Pluronic F-127, 1 mg/mL α-casein, and 20 μM Taxol) and incubated for 1 hour to fully block the glass surface. The slide chamber was then washed twice with 20 μL of BB, and the solution inside the chamber was completely removed. Subsequently, 12 μL of 1 mg/mL streptavidin was flowed into the chamber and incubated for 10 minutes. The chamber was then washed three times with 20 μL of BB. Finally, 20 μL of BB that contained the suspended MTs and 10 μM paclitaxel was flowed into the chamber and immediately washed with 40 μL of BB. Polarity-marked MTs were prepared as described previously^38^.

### Single-molecule motility assay

To dimerize the single-headed dynein constructs, Cy3-labeled anti-GFP antibodies were used as previously described^38^. Dynein constructs were diluted to the appropriate concentration using HME30 (30 mM HEPES, 2 mM MgCl_2_, and 1 mM EGTA) and incubated on ice with Cy3-labeled, anti-GFP antibodies for 10 minutes. A final motility buffer (MB) containing 1 mM ATP, 1 mg/mL α-casein, 20 μM Taxol, 75 mM KCl, 2 mM Trolox, Gloxy, 1 mM TCEP, and the motor-antibody mixture was flowed into the chamber. MTs and dynein were visualized with a custom-built TIRF microscope equipped with an Andor iXon Ultra EMCCD. MTs were first imaged by taking a single-frame snapshot. Dynein was then imaged with an acquisition time of 500 ms and a total 500 frames were acquired for each movie. Finally, the MTs were imaged again by taking a snapshot to check for stage drift. Movies with significant drift were not analyzed. Each sample was imaged no longer than 15 minutes. The kymographs were generated using Fiji and the data were analyzed using a custom written MATLAB program.

### Unbinding-force assay

Unbinding-force measurements were performed using a custom-built force-fluorescence inverted microscopy as described previously^83^. Briefly, polarity-marked MTs were attached to the flow chamber coverslip as described above. Yeast dynein was diluted to the appropriate concentration using trapping buffer (30 mM HEPES, 2 mM MgAc_2_, 1 mM EGTA, 20 μM paclitaxel, 20 mM glucose, and 2 mM Trolox, pH 7.2) and incubated on ice with anti-GFP antibody Fab fragment-coated ∼1-μm diameter beads (980 nm, carboxyl-modified polystyrene microspheres, Bangs Laboratories) for 10 minutes. To remove free unbound motors, the beads were centrifuged at 4°C for 2 minutes at 3000 × g, followed by supernatant removal. The beads were then resuspended with 40 μL trapping buffer containing 0.75 mg/mL α-casein, Gloxy (a glucose oxidase and catalase-based oxygen scavenging system) and 6.6 units/mL Apyrase (to remove residual ATP for the apo state experiments) or 1 mM ATP. A bead bearing a dynein MD was then held in an optical trap as the nano-positioning stage swept back and forth parallel to a surface-bound MT. The stage speed was adjusted to produce an apparent loading rate of 5.6 pN/s (the true loading rate depends on the compliance of the motor and is smaller than the apparent rate^38^).

### Analysis of data generated by the constant-pulling assay

As we showed previously^58^, the largest forces in our constant-pulling unbinding experiments (**Fig. 3a**) usually occur when a bead rebinds to the MT before returning to the trap center. We call these secondary binding/unbinding events. For primary events, zero force was applied to the MD immediately after MT binding (F_start_ = 0), but for secondary events, F_start_ > 0. Because the history of force applied to the bond depends on F_start_, we only focused on primary events (**Fig. 3a**, right). Measurements from multiple beads and experiments under the same conditions were pooled together and used to generate unbinding-force histograms with 1-pN bins (as our force detection limit is ∼0.3 pN, we only plotted the force-dependent unbinding rates for forces above 0.5 pN). To compare the unbinding-force distributions and the derived force-dependent unbinding rates for both loading directions, we plotted the data as a function of the absolute force values. Normalized histograms that approximated the probability density functions for unbinding at a given force were then calculated. The value of each bin was divided by *N*, the total number of unbinding-force measurements. Because the unbinding-force distributions were not normally distributed, we estimated the sampling error by bootstrapping. For each histogram, 95% confidence intervals (CIs) for the mean statistic were calculated using the MATLAB bootci() function as described previously^58^. To estimate *p*-values when comparing means of different distributions, we first created a dataset representing the sampling distribution of the mean for each original dataset. First, we bootstrapped 10^5^ means with the MATLAB function bootstrp(). We then subtracted these means pairwise to create a dataset representing the sampling distribution of the difference of the means. From each measurement in this dataset, we subtracted the mean difference of the means, which shifted the mean of the distribution to zero. This transformation was consistent with the null hypothesis of no difference between the original unbinding-force distribution means. The *p*-value (*p*_*m*_) was then calculated as the proportion of the bootstrapped mean differences that were greater than or equal to the difference observed between the means of the original datasets (two-tailed test). Similar to our recent work^38^, we considered the compliance of the dynein motor and bead linkage, which resulted in a force-dependent loading rate. Uncertainty arises in the calculated unbinding rates as a function of force due to limited statistics for larger forces. To reduce this uncertainty, we used a kernel density estimator to describe the probability density functions of the measured unbinding forces before transforming them into force-dependent unbinding rates^84^.

## Acknowledgements

All authors were supported by the National Institute of Health (NIH) grant R01GM098469. X. L. also received support from a Pilot Project Grant from the Rose F. Kennedy Intellectual and Developmental Disabilities Research Center at the Albert Einstein College of Medicine.

## Author contributions

X.L., L.R., and A.G. designed the research. X.L. performed the research and engineered the mutant yeast strains. X.L. also produced and purified the proteins. X.L. and A.G. analyzed the data and wrote the paper.

## Notes

### Competing Interest Statement

The authors have declared no competing interest.

